# A reference genome and transcriptome of haustorial development in *Pedicularis groenlandica* reveal diverse trajectories of haustoria-associated gene evolution in parasitic plants

**DOI:** 10.1101/2025.11.20.689508

**Authors:** Rachel O. Cohen, Anri Chomentowska, Liming Cai, Deren A. R. Eaton

## Abstract

- Novel traits frequently evolve by co-opting existing genetic pathways through direct repurposing of existing genes or neofunctionalization of duplicated genes. Parasitism represents a major innovation in plants, where a novel organ—the haustorium—evolved to penetrate hosts and extract water and nutrients. Previous studies hypothesized that haustoria-associated genes primarily evolve from root and pollen-associated pathways.
- To examine evolutionary trajectories of haustoria-associated genes, we generated a chromosome-scale genome of Pedicularis groenlandica (Orobanchaceae) and sampled transcriptomes throughout haustorial development. We examined differential expression of haustoria-associated genes and their paralogs and investigated orthology among haustoria-associated genes in five parasitic plants.
- We identified 5,635 haustoria-associated genes in P. groenlandica, of which a greater proportion were associated with pollen tubes (67%) than roots (33%), evidenced by being differentially expressed in pollen tubes, nested within pollen tube-associated gene families, or both. Haustoria-associated genes with paralogs that arose after the evolution of parasitism in Orobanchaceae are more likely to be uniquely expressed in haustoria, consistent with neofunctionalization.
- Our results support both pleiotropy and neofunctionalization as mechanisms by which genetic pathways are co-opted for haustorial function. Haustoria-associated genes are highly lineage-specific, highlighting a dynamic and ongoing process of haustorial co-option of genes among parasitic plant lineages.

## Introduction

Approximately 1% of flowering plants are parasites on other plants, a strategy that has evolved independently around a dozen times in angiosperms (Westwood *et al*., 2010). Parasitic plants use a novel organ, the haustorium, to invade host tissues and establish a vascular connection to extract water and/or nutrients. The repeated convergent evolution of haustoria—including their origination from both root (e.g., *Striga*) and stem tissue (e.g., *Cuscuta*) in different lineages— raises fundamental questions about the genomic, developmental, and ecological precursors that enable this transition. Comparative studies have revealed extensive anatomical and physiological convergence in haustoria across a range of lineages (Fernández-Aparicio *et al*., 2016; Yoshida *et al*., 2016; Masumoto *et al*., 2021) and recent genomic studies have identified both lineage-specific and shared molecular pathways associated with haustorial development and function (Westwood *et al*., 2012; Zhang *et al*., 2015; Jhu *et al*., 2021; Kokla *et al*., 2021; Bawin *et al*., 2022, 2024). Despite our expanding knowledge of the genes that govern haustorial attachment across parasitic lineages, the evolutionary processes by which genes have been co-opted for haustorial function remain poorly understood.

Parasitic plants span a continuum of host dependence based on developmental requirements (facultative versus obligate parasites), and trophic capacity (hemiparasites, which retain photosynthetic capacity, versus holoparasites, which are fully heterotrophic). Facultative hemiparasites produce lateral haustoria that connect primarily to host xylem, whereas obligate parasites form a terminal haustorium that connects to both xylem and phloem (Press & Graves, 1995; Yoshida *et al*., 2016). It has been hypothesized that parasitism evolves in a stepwise fashion, with facultative hemiparasitism representing an initial stage that in some lineages gave rise to obligate hemiparasites, and, ultimately, holoparasites (Westwood et al., 2010; Xu *et al*., 2022). The family Orobanchaceae uniquely spans this full spectrum (including non-parasitic taxa), and encompasses economically significant weeds such as witchweeds (*Striga*) and broomrapes (*Orobanche*) that cause extensive damage to crops worldwide (Joel *et al*., 2013; Li *et al*., 2019). As a result, Orobanchaceae has emerged as a model system for the study of haustorial physiology and the genomic basis of parasitic plant evolution.

Haustoria appear, at least in part, to co-opt genetic pathways active in other tissues. Indeed, plants frequently redeploy existing gene repertoires to acquire novel functions and life-history strategies. For example, the co-option of MADS-box genes for floral organ development (originally functioning in gametophyte-sporophyte transitions; Niklas & Kutschera, 2009) and terpene synthases for specialized volatile production and herbivore defense (originally functioning in basic metabolic processes; Abbas et al., 2017). It has been hypothesized that Orobanchaceae, being root parasites, predominantly co-opt root-associated genetic pathways for haustorial function, in addition to some non-root-associated genetic pathways. Consistent with this view, Yang et al. (2015) found that orthologs of haustoria-associated genes are frequently highly expressed in roots of non-parasitic plant species. Curiously, they additionally found that haustorial orthologs are often differentially expressed in pollen and flowers, a result replicated by Yoshida *et al*. (2019). We hypothesize that the relationship between haustoria- and pollen-associated genes results from shared functions between haustoria and pollen tubes, including signal detection, polarized tip growth, cell-wall remodeling, and traversal of foreign tissues. The discovery of haustoria-associated gene homologs expressed in other tissue types may suggest that neofunctionalization of paralogs gave rise to haustorial function. Alternatively, these genes may participate pleiotropically in haustorial development while still functioning in other tissues through tissue-specific regulation.

Novel genes required for haustorial development have been predominantly hypothesized to arise following whole-genome duplications (WGDs), which resulted in duplicate copies that could be co-opted and neofunctionalized (Vogel *et al*., 2018; Lyko & Wicke, 2021). Signals of ancient WGDs are indeed detectable in Orobanchaceae (Yang *et al*., 2015; Xu *et al*., 2022) and *Cuscuta* (Sun *et al*., 2018), a distantly-related stem-parasitic lineage. Yang et al. (2015) reported that more than half of gene families containing haustoria-associated genes in three species of Orobanchaceae show evidence of ancestral gene duplications, which they hypothesize to be the result of WGDs. In both Orobanchaceae and *Cuscuta*, the WGDs most plausibly linked to haustorial evolution are shared with non-parasitic relatives. Furthermore, Yang et al. (2015) did not evaluate alternative evolutionary hypotheses in addition to WGDs that could explain how genes are co-opted for haustorial function, like other classes of gene duplications (*i.e.* tandem duplications) or pleiotropy. Although gene duplication is a well-recognized engine of evolutionary novelty (Ohno, 1970), plants tend to experience gene and genome duplications at high rates (Moore & Purugganan, 2005) and these phenomena cannot be linked to parasitic plant evolution specifically. Thus, while some haustorial genes may trace to WGD-derived neofunctionalization, additional evidence is needed to evaluate diverse evolutionary explanations for the co-option of genes for haustorial function following transitions to parasitism.

Over the past decade, newly developed genetic resources in Orobanchaceae have created the opportunity to investigate gene family evolution and tissue co-expression patterns of haustoria-associated genes within a comparative framework (Westwood *et al*., 2012; Yang *et al*., 2015; Xu *et al*., 2022). Many of these studies (Westwood et al. 2012, Yang et al. 2015, Yoshida et al. 2019) leveraged the same transcriptome datasets from the Parasitic Plant Genome Project (Westwood *et al*., 2012) and contig-level genome assemblies. Advances in genome and transcriptome sequencing and assembly now allow for more accurate identification of haustoria-associated expression, orthology inference, and reconstruction of gene regulatory modules, all of which will aid our ability to determine the identity and function of haustoria-associated genes and evaluate diverse evolutionary hypotheses for their origins.

In this study, we assembled a new chromosome-scale reference genome and annotation of the facultative hemiparasite *Pedicularis groenlandica* (Orobanchaceae), and generated a transcriptomic time series of haustorial development, representing only the third such haustorial expression dataset for a high-quality chromosome-scale genome. Using these data we: (1) identify haustoria-associated genes in *P. groenlandica* and evaluate their chromosomal distribution and overlap in expression with other tissue types; (2) infer co-expression modules and their tissue overlap, and predict regulatory motifs associated with haustorial expression; (3) test the role of neofunctionalization in haustoria-associated gene evolution by comparing tissue-specific expression between paralogs that diverged before versus after the evolution of parasitism in Orobanchaceae; and (4) identify orthology among haustoria-associated genes in *P. groenlandica* and other parasitic plant lineages.

## Materials and Methods

### Study system

*Pedicularis groenlandica* (Elephant’s head lousewort) is an iconic flowering plant of western North American alpine meadows. *Pedicularis* are facultative hemiparasites that attach to a range of host plants through lateral haustoria, most often parasitizing sedges, grasses, and legumes (Piehl, 1963; Ren *et al*., 2010; DiGiovanni *et al*., 2017). *Pedicularis* is well-studied for its distinctive pollination biology (Macior, 1968; Grant, 1994; Eaton *et al*., 2012; Tong & Huang, 2016; Cohen *et al*., 2025), but little attention has been paid to parasitism within the genus. Two chromosome-scale *Pedicularis* genome assemblies have been published: *P. cranolopha* (Xu *et al*. 2022) and *P. kansuensis* (Fang *et al*., 2024). Both studies considered how parasitism has impacted *Pedicularis* genome evolution, including investigations of gene loss, transposable elements, and horizontal gene transfer, but neither examined haustorial gene expression.

### Tissue collection and genome sequencing

Plant tissues for genome sequencing of *P. groenlandica* were collected at Rocky Mountain Biological Laboratory in Gothic, Colorado, USA in the summer of 2024. Flow cytometry was performed on fresh leaf tissue to estimate genome size, and 5.0g of leaf tissue was sent to Cold Spring Harbor Laboratory (Cold Spring Harbor, NY, USA) for high-molecular weight DNA extraction, PacBio Hifi library preparation and sequencing on a PacBio Revio, and HiC library preparation and sequencing on an Illumina NextSeq2000 to generate 2 x 150bp reads.

For genome annotation and differential expression studies, we collected three biological replicates of young leaf, stem, flower bud, pollen tube, root, and mature haustoria tissues into RNAlater (Invitrogen), which was stored at -80°C until extraction. To sample pollen tubes, we germinated fresh *P. groenlandica* pollen on pollen tube growth media, using the methodology employed in Cohen *et al*. (2025). After 24 hours, germinated pollen tubes were flash frozen over a dry ice-ethanol slurry and stored at -80°C until extraction. RNA was extracted from all tissues using the Qiagen RNeasy Plant Minikit, and Illumina libraries were prepared and sequenced via Illumina MiSeq at Admera Health LLC (South Plainsfield, NJ, USA) to generate 2 x 150bp reads.

### Genome Assembly and Annotation

PacBio Hifi reads were filtered to retain only sequences >10kb in length, and assembled into a phased contig-level assembly using Hifiasm v0.19.8 run in “HiC mode” (Cheng *et al*., 2021). The primary contig-level assembly was analyzed with the Arima Genomics Mapping Pipeline v.03 using YAHS for scaffolding (Zhou *et al*., 2023), juicer (Durand *et al*., 2016) to convert the scaffolded assembly into a Hi-C contact map, and JuiceBox Assembly Tools (JBAT; Durand *et al*., 2016) for manual curation. We evaluated the completeness of our genome with BUSCO v.5.7.0 against eudicots_odb10.1 (Seppey *et al*., 2019), then soft-masked the genome using RepeatModeler v2.0.6 (Flynn *et al*., 2020) followed by RepeatMasker v4.1.7p1 (Smit, Hubley & Green 2013–2015).

We structurally annotated the genome using BRAKER3 v3.0.8 with both RNAseq and protein evidence (Stanke *et al*., 2006, 2008; Quinlan, 2014; Kovaka *et al*., 2019; Pertea & Pertea, 2020; Brůna *et al*., 2024; Gabriel *et al*., 2024). We used RNAseq from all tissues and protein evidence from the Viridiplantae dataset in ORTHODB v.10 (Zdobnov *et al*., 2021), which were trimmed of adapters and low-quality bases (<Q30) using TrimGalore v0.6.10 (Krueger, 2015), and quality assessed using FastQC v0.12.1 (Andrews, 2010). The annotation completeness was evaluated with BUSCO v.5.7.0 against the eudicots_odb10.1 dataset in transcriptome mode. We functionally annotated the genome using InterProScan with the –goterms tag, and identified the closest *Arabidopsis thaliana* ortholog of each gene with BLAST v2.16.0 (Ye *et al*., 2006). We conducted a synteny analysis in order to compare *P. groenlandica* to *P. cranolopha* and *O. cumana* (Xu *et al*., 2022) using MCScan (JCVItools; Tang *et al*., 2024).

### Haustorial time series tissue sampling and sequencing

To generate a time series of gene expression during haustorial growth and development we sampled RNA from *P. groenlandica* haustoria grown in petri dishes while interacting with a host plant. Seeds for this assay were collected from mature seed pods of *P. groenlandica* in the same population as the genome plant, removed from their pods, cleaned of excess tissue, and stored in dry envelopes at 4°C. *Pedicularis groenlandica* do not require hosts for germination and early growth. We germinated parasites and hosts separately, following the protocol of Westwood et al. (2012) for staged haustorial tissue collection.

To promote germination, seeds of *P. groenlandica* were treated with a 200ppM solution of gibberellic acid for 15 minutes, and then sterilized in a solution of 30% bleach, 10% ethanol, 60% sterile de-ionized water, and 1 drop of triton, by submerging seeds in the sterilizing solution with agitation for 5 minutes, followed by three washes of shaking in sterile water for three minutes each, and then dried completely. We germinated seeds on plates with 10% Murashige-Skoog (MS) media solution prepared in an autoclave. Seeds were placed ∼2cm apart on each plate, and plates were wrapped in Parafilm and placed in a growth chamber on a diurnal cycle with 18 hours of light at 22°C and 6 hours of dark at 18°C.

*Arabidopsis thaliana* was chosen as the host plant in our haustorial assays due to its fast growth rate and because it has served as a successful host in previous *Pedicularis* haustorial assays (Kee Jia Yee, *personal communication*). Seeds of *A. thaliana* accession Col-0/Rédei-L209478*A* were purchased from Lehle seeds (Round Rock, Texas). Due to their faster rate of germination, we began germinating *A. thaliana* seeds three days after *P. groenlandica* germination, in separate plates, following the same germination protocol. Seeds were monitored for germination over a 10 day period, and only those with both cotyledons open and primary root growth >2cm were selected for assays. To initiate the assay, seedlings of *P. groenlandica* and *A. thaliana* were carefully removed using sterilized forceps and placed together on a single, sterile, 10% MS plate. We used nutrient-free media at this stage to promote haustorial development. We initially implemented trials in which the *A. thaliana* root was wrapped around the *P. groenlandica* root, but found that placing it ∼2mm away from *P. groenlandica* promoted greater root and lateral haustorial growth towards *A. thaliana*, and so we proceeded to conduct all trials using this methodology.

Our assay included tissue sampled for RNA extraction at four distinct stages (1-4) to establish a time series of haustorial development and attachment (**Fig. 1**). We additionally sampled 1 cm of root tissue from *P. groenlandica* prior to host exposure (stage 0). In the first treatment step (stage 1), we sampled 1 cm of root tissue 24 hours after exposure to the host plant. At the second treatment step (stage 2), bulbous “swelling” haustoria were observed, which form prior to host penetration and attachment. We sampled haustoria, including approximately 1mm of root tissue on each side of the haustorium. In the final treatment step (stage 3), haustoria have fully penetrated and attached to the host. We confirmed this by pulling lightly on the *A. thaliana* root, if the *P. groenlandica* root moved with it we considered the haustoria fully attached. The haustoria were again sampled including 1mm of *P. groenlandica* root tissue above and below. Finally, our assay also included mature, field-collected haustoria (stage 4) sampled from plants dug up during tissue collection for genome annotation. All tissues were collected in triplicate, immediately flash-frozen in liquid nitrogen and stored at -80°C until extraction. RNA was extracted using the Qiagen RNeasy Plant Minikit, and libraries were prepared using Poly-A selection and sequenced via Illumina MiSeq at Admera Health LLC (South Plainsfield, NJ, USA) to generate 2 x 150bp reads.

**Fig. 1.**
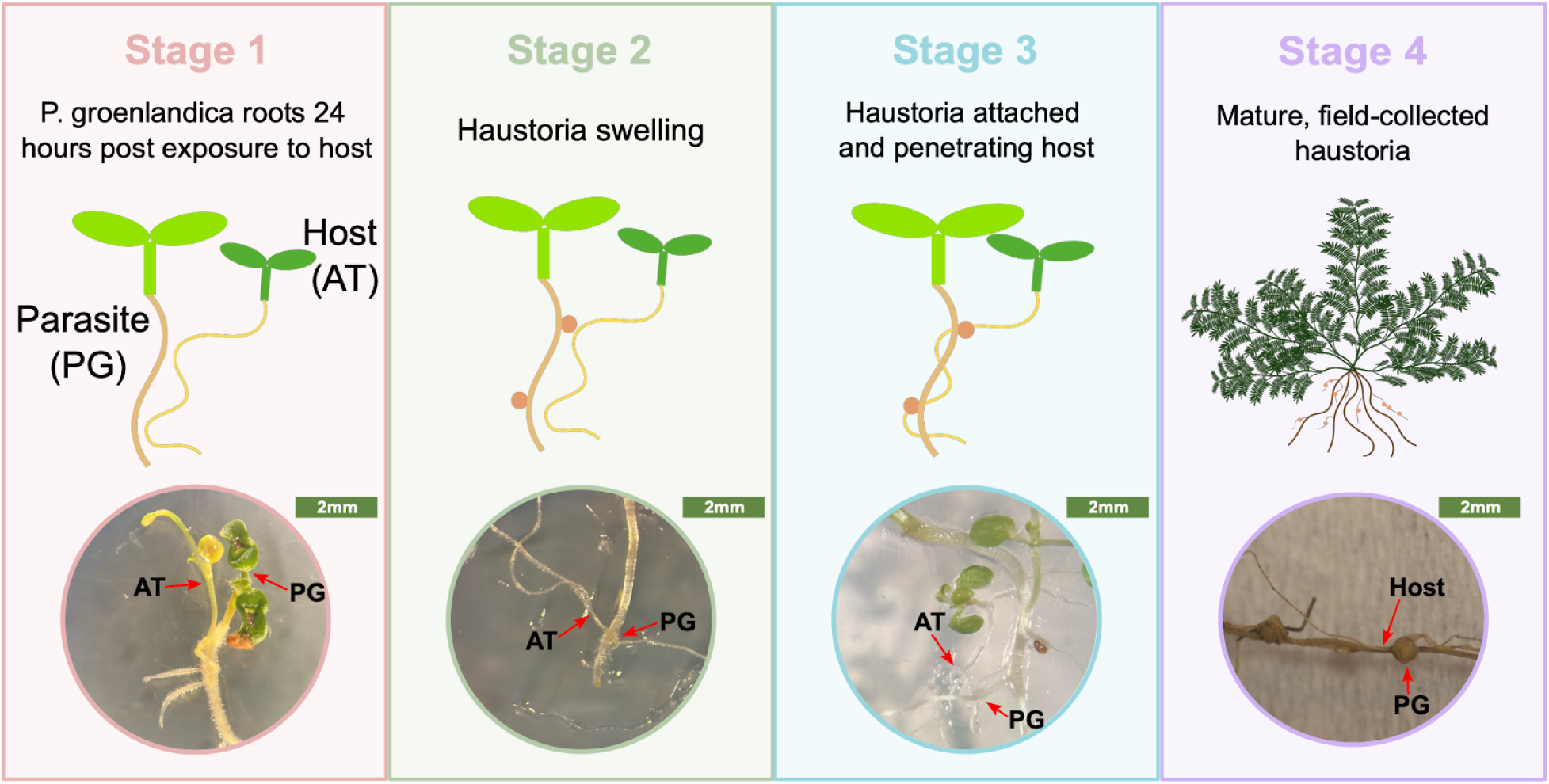
Stages of haustorial development and attachment sampled in an assay involving the facultative hemiparasite *Pedicularis groenlandica* (PG). Stages 1-3 were sampled from haustoria attachment assays conducted in petri dishes, using *Arabidopsis thaliana* (AT) as a host plant. For stage 4, we sampled mature haustoria in the field, where *P. groenlandica* was attached to a natural *Medicago* host (labeled “Host”). Sampling stages were selected following Westwood et al. (2012).

### Differential expression analysis and gene network coexpression analysis in haustorial tissues

As described for the annotation RNA sequences, RNAseq reads were trimmed of adapters and low-quality bases (<Q30) using TrimGalore v0.6.10 (Krueger, 2015), with quality further assessed by FastQC v0.12.1 (Andrews, 2010). All filtered reads, including from leaf, stem, root, flower bud, and pollen tubes, were mapped to the final *P. groenlandica* genome using STAR v2.7.11b (Dobin *et al*., 2013). Read counts were obtained individually for each replicate at each stage of the haustorial assay (0-4), as well as for each non-parasitic tissue. Following alignment, we used featureCounts (Liao *et al*., 2014) to count reads mapped to each annotated gene.

To identify differentially expressed genes, we analyzed expression counts in the R package DESeq2 v1.42.1 (Love *et al*., 2014). We compared haustorial growth stages (Stages 1-4) against non-parasitic tissue-types (leaf, stem, root, flower bud, pollen tubes and Stage 0) to identify genes that were either up- or down-regulated in each haustoria time stage compared to non-parasitic tissue. Prior to running DESeq2, we excluded low count genes (counts <10). To visually confirm the expected similarity among replicates within tissues, and between more similar tissues, we performed a principal components analysis of the gene expression data from DESeq2. A gene was determined to be differentially expressed in haustoria if it had a log2FoldChange>2 (up-regulated) or <–2 (down-regulated) relative to non-parasitic tissues, and an adjusted p-value <0.05. We compiled a list of “haustoria-associated” genes, which includes the set of genes that were differentially expressed in at least one stage (1-4) of the haustorial assay. We assessed the distribution of haustoria-associated genes across the *P. groenlandica* genome and calculated the correlation between haustoria-associated gene density and overall gene density using Pearson’s correlation coefficient and a Student’s T-test.

To identify modules of co-expressed genes, we implemented weighted gene co-expression network analysis (WGCNA). Following the methodology of Gilman *et al*. (2022), we input the quantified transcripts per million (TPM) for each gene to WGCNA v1.73 in R (Langfelder & Horvath, 2008). We assessed which of the resultant co-expression gene modules were correlated with gene expression in each of the tissues included in analyses (haustoria, leaf, stem, root, flower bud, pollen tubes) by calculating the Pearson’s correlation coefficient and assessing significance by a student’s asymptotic t-test. A module was designated as haustoria-associated if its expression was significantly correlated with haustorial samples (stages 1-4) compared to non-haustorial samples (stage 0 and other tissues). To determine functional descriptions of the genes in each module, we first identified their orthologs in *A. thaliana* through BLAST against the TAIR *A. thaliana* Genome V (Lamesch *et al*., 2012), then determined the Gene Ontology biological process terms that were enriched in each module using the “enrichGO” function in the *clusterProfiler* in R (v4.10.1; Wu *et al*., 2021).

### Overlap of differentially expressed genes and regulatory motifs in haustoria and non-parasitic tissues

To investigate the hypothesis that haustoria-associated genes may also be differentially expressed in other tissues, we measured the overlap in genes that were differentially expressed in both haustoria and leaf, stem, root, flower bud, pollen tubes. Differential expression analysis requires a pairwise comparison (*i.e.* genes are up-regulated in haustoria *compared* to pollen tubes). We independently quantified differential expression in each individual overlap analysis, making sure haustoria and the focal tissue were always compared to the same ‘control’ set. For example, when investigating haustoria- and pollen tube-associated gene overlap, we identified genes differentially expressed in haustoria compared to leaf, stem, root, and flower bud, and genes differentially expressed in pollen tubes compared to the same set of tissues. We proceeded in this manner, systematically comparing haustoria and the focal tissue to the identical control sets, then assessed how many of the genes overlapped among haustoria and the focal tissue. Through this analysis, we identified not only the overlap in differential expression of haustoria-associated genes with other tissues, but also identified a set of “haustoria-specific” genes that are only differentially expressed in haustoria.

We additionally investigated whether co-expressed networks of haustorial-associated genes shared enriched regulatory motifs with other tissue-associated modules. To identify enriched motifs associated with each tissue, we first identified WGCNA modules of genes whose expression patterns were correlated with each tissue, using Pearson’s correlation coefficient, as above. Because WGCNA modules contain genes that are not tissue-associated as determined by the differential expression analysis, we searched for enriched motifs within 2kb of only tissue-associated genes in each tissue-associated module using XSTREME (Meme v5.5.8; Bailey *et al*., 2009) following methods in Chomentowska *et al*. (2025). We identified enriched motifs that overlapped in haustoria and other tissues in both sequence and locus. Similar to our approach for identifying overlap in differentially expressed genes among haustoria and non-parasitic tissues, for each individual motif overlap analysis, we independently identified coexpression modules with WGCNA and evaluated module correlation with each focal tissue excluding haustoria, then haustoria modules excluding the focal tissue such that the haustoria and focal tissue were again compared to an identical group of “control” tissues when determining module-tissue correlation.

Because pollen tubes and haustoria had the highest overlap in terms of tissue-associated genes and regulatory motifs, they were further evaluated. For the set of genes that overlapped between haustoria and pollen tubes we assigned molecular function GO terms by identifying their orthologs in *A. thaliana*, then using the “enrichGO” function in the R package *clusterProfiler* (v4.10.1; Wu et al., 2021). Using the modules constructed during overlap analyses, we asked if haustoria- and pollen tube-associated genes were similarly grouped into co-expression modules more often than expected by chance. To do this, we constructed a contingency matrix which included the ‘observed values’ of overlapping haustoria- and pollen tube-associated genes assigned to haustoria- and pollen tube-associated modules, respectively. To set null expectations, we performed a permutation-based test in which we randomly shuffled module assignments while preserving the number of genes in each module. We conducted 2,000 permutations and recalculated the overlap contingency table with these ‘expected values’, then recorded the number of times observed values were greater than or equal to expected values and calculated empirical *p*-values. To account for multiple testing across all module pairs, *p*-values were adjusted using Benjamini–Hochberg false discovery rate (FDR). Finally, we functionally annotated the shared enriched motifs by identifying the family of transcription factors with which each motif interacts using TOMTOM (Meme v5.5.8; Bailey et al., 2009), comparing our enriched motifs against known motif annotations in the JASPAR database (Rauluseviciute *et al*., 2024).

### Analysis of haustorial gene trees and gene duplications

To evaluate the hypothesis that haustoria-associated genes evolved by neofunctionalization following gene or genome duplication events in a non-parasitic ancestor of Orobanchaceae, we examined the phylogenetic relationships of haustoria-associated genes with respect to their paralogs and orthologs, and their differences in tissue-specific expression. We downloaded protein sequences of four publicly available non-parasitic genomes from NCBI Genbank and JGI Phytozome: *Mimulus guttatus* (NONTOL v4.0, DOE-JGI), *Salvia miltiorrhiza* (GCF_028751815.1; Pan *et al*., 2023), *Solanum lycopersicum* (ITAG5.0; Zhou *et al*., 2022), and *Arabidopsis thaliana* (TAIR10; Lamesch *et al*., 2012), and assembled orthogroups using Orthofinder v3.0.8 (Emms & Kelly, 2019) run with default settings. Orthogroups were filtered to retain only those that contained one or more haustoria-associated genes. Multiple sequence alignments from Orthofinder were cleaned and trimmed using a custom Python script, and gene trees were inferred by RAXML-ng v1.2.2 (Kozlov *et al*., 2019) with support assessed from 100 non-parametric bootstraps. Many trees contain multiple gene copies per species, so we used the Duplication-Transfer-Loss model in GeneRax v2.1.3 (Morel *et al*., 2020) to obtain reconciled and rooted genes trees based on the species tree topology, retaining trees with >3 genes (*i.e.* tips), according to default GeneRax settings.

For each *P. groenlandica* haustoria-associated gene in each rooted orthogroup tree, we identified paralogs and recorded their tissue-specific expression. Tree-based operations were performed in the Python package toytree v.3.0.10 (Eaton, 2020) to identify: the closest non-haustoria-associated paralog that is more closely related than the closest ortholog (i.e., it arose by duplication within Orobanchaceae); and the closest non-haustoria-associated paralog that is equal to or more distantly related than the closest ortholog (*i.e.*, it arose by duplication in an ancestor of Orobanchaceae; *see* ***Fig. 5a***). If multiple paralogs were equally related to the haustoria-associated gene, one was chosen at random. For each gene, we recorded the tissue in which it was differentially expressed, as measured above.

To validate our approach, we visually examined many gene trees, including for several selected genes with known haustorial function. We searched TAIR10 (Lamesch et al., 2012) to identify the *A. thaliana* gene locus identifiers for homologs of genes that have been experimentally shown to affect haustoria development and hormone signaling or pollen tube growth and signaling. This includes ROOT MERISTEM GROWTH FACTOR (RGF), which (Fishman *et al*., 2025)demonstrated is necessary for haustorial development in the hemiparasite *Phtheirospermum japonicum*; KARRAKIN INSENSITIVE-2 (KAI2), which has been shown to have paralogs that enable ultra-sensitive strigalactone receptors in Orobanchaceae (Wang *et al*., 2021; Takei *et al*., 2024), two deep homologs of KAI2, DWARF14 (D14), and DWARF14-LIKE2 (DLK2), and pollen tube growth and guidance genes Cyclic Nucleotide-Gated Channel 18 (CNGC18; von Besser et al., 2006), HAPLESS 2 (HAP2; Daher & Geitmann, 2012), and Actin-Depolymerizing Factor 7 (ADF7; Gao *et al*., 2016).

### Modeling the effect of gene duplications

To quantify the relative importance of gene duplications to the evolution of haustorial specificity in gene expression, we compared the number of genes that are haustoria-specific (*i.e.*, only differentially expressed in haustoria) to those that are differentially expressed in multiple tissues including haustoria, and whether haustoria-specific expression is predicted by the orthogroup having recent and/or ancient duplications. For each haustoria-associated gene, we recorded whether it was haustoria-specific, whether a recent paralog exists in the gene tree, and whether an ancient paralog exists. The number of *P. groenlandica* genes in each orthogroup was included as a covariate (log transformed and centered before fitting). We fit a population-averaged logistic model using generalized estimating equations (GEE; Binomial, logit link) with genes clustered by orthogroup, an exchangeable working correlation, and robust (sandwich) standard errors. Models were fit using statsmodels (Seabold & Perktold, 2010) in Python. Two-sided Wald tests were used to assess coefficients, with odds ratios and 95% confidence intervals. We compared the effect of recent versus ancient paralogs via a Wald X^2^ test, and computed average marginal effects.

### Orthology and expression overlap of haustoria-associated genes among parasitic plant species

To investigate the extent to which haustoria-associated genes in *P. groenlandica* are shared with other parasitic plant lineages, we examined differential expression and orthology with other haustoria development datasets. We restricted our analyses to four datasets with transcriptomic time-series of haustorial development analogous to our dataset (*i.e.*, RNA sampled from at least stages 1-3). This included: *Cuscuta campestris* (Bawin *et al*., 2024), *Phtheirospermum japonicum* (Kokla *et al*., 2021), *Striga hermonthica* (Westwood *et al*., 2012), and *Orobanche aegyptiaca* (Westwood *et al*., 2012). The quality of these datasets varied—*S. hermonthica* and *O. aegyptiaca* were generated in 2012, whereas *C. campestris* and *P. japonicum* were generated using more modern technologies since 2020. We utilized the control haustorial dataset from Kokla et al. (2022) whose main research focus was repression of haustorial formation by nitrogen, rather than the process of haustorial formation itself.

In order to identify haustoria-associated genes by stage in each of these datasets, we followed the same pipeline that was used in *P. groenlandica*. A publicly available reference genome is available for each species with the exception of *O. aegyptiaca*. For this taxon, we instead mapped reads to the *O. cumana var. cumana* genome (Xu et al., 2022). Because reads mapped at a sufficiently high rate (average 66.8% mapping rate across all tissues and stages), we proceeded with the remainder of the pipeline. Genes from *P. groenlandica* and the four published genomes were classified into orthogroups using Orthofinder v3.0.8 (Emms & Kelly, 2019). We quantified orthogroup overlap among species’ haustoria-associated genes, then functionally annotated the haustorial orthologs by identifying their *A. thaliana* orthologs, followed by conducting a Gene Ontology term enrichment analysis using *clusterProfiler*, as described above.

## Results

### The P. groenlandica genome

The *P. groenlandica* genome represents a new, high-quality genomic resource in *Pedicularis* and Orobanchaceae. The assembled haploid genome size is 1.38Gb, consistent with the flow cytometry estimate (C=1.59; **Fig. S1**). We scaffolded the genome into eight highly contiguous chromosomes which include 91.6% of nucleotide base pairs (**Fig. 2**). The BUSCO completeness score was 96.5% (Single-copy: 90.5%, Duplicates: 6.1%). The *P. groenlandica* genome contained a high proportion of repetitive elements, with repeats making up 76.48% of the genome. A large proportion of repeats (44.14%) are long terminal repeats (LTRs). Our genome has high synteny to *P. cranolopha* (Xu *et al*., 2022), but exhibit rampant rearrangement with respect to *Orobanche cumana var. Cumana*. We annotated 27,006 unique protein coding genes. The BUSCO score of the annotation was 94.3% (Single-copy: 59.6%, Duplicates: 34.7%).

**Fig. 2.**
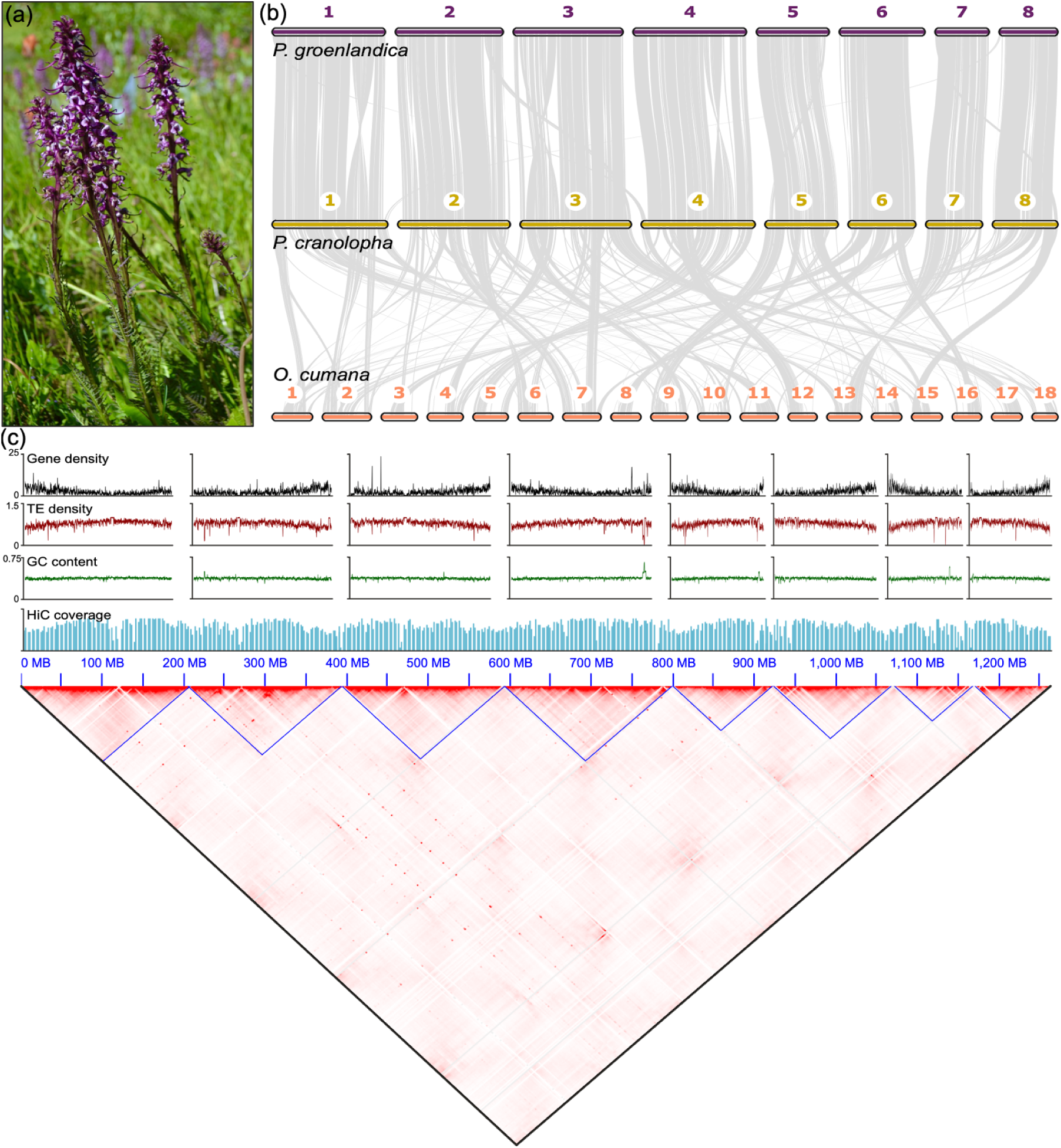
Chromosome-scale assembly of the *Pedicularis groenlandica* genome. **a)** Photo of *P. groenlandica* (Elephant’s head lousewort). **b)** Gene synteny between the eight chromosomes of *P. groenlandica* and *P. cranolopha*. Both *Pedicularis* species have highly rearranged genomes with respect to the *Orobanche cumana var. cumana* genome (Xu *et al*., 2022). **c)** Gene density, transposable element (TE) density, GC content, and HiC sequence coverage statistics along 10kb sliding windows in the *P. groenlandica* genome. Genes and TE’s are distributed evenly across the genome, and there are few peaks in GC content. We show the HiC contact-map shows 8 contiguous chromosomes in *P. groenlandica*.during correlation tests. 2,047 genes were differentially expressed, showing enrichment for light response and photosynthesis biological process GO terms.

#### Haustoria-associated gene expression, function, and regulation

Across our RNAseq libraries, we obtained an average of 27.84 million reads per replicate per tissue, of which an median of 20.89 million reads (83.01%) uniquely mapped to the *P. groenlandica* genome. All protein coding genes were represented in the final count dataset. Differential expression analysis in DESeq2 identified 5,635 haustoria-associated genes across all four stages (**Fig. 3a-b**). The density of haustoria-associated genes across chromosomes was highly correlated with the density of all genes (P<0.001; **Fig. 3d**).

**Fig. 3.**
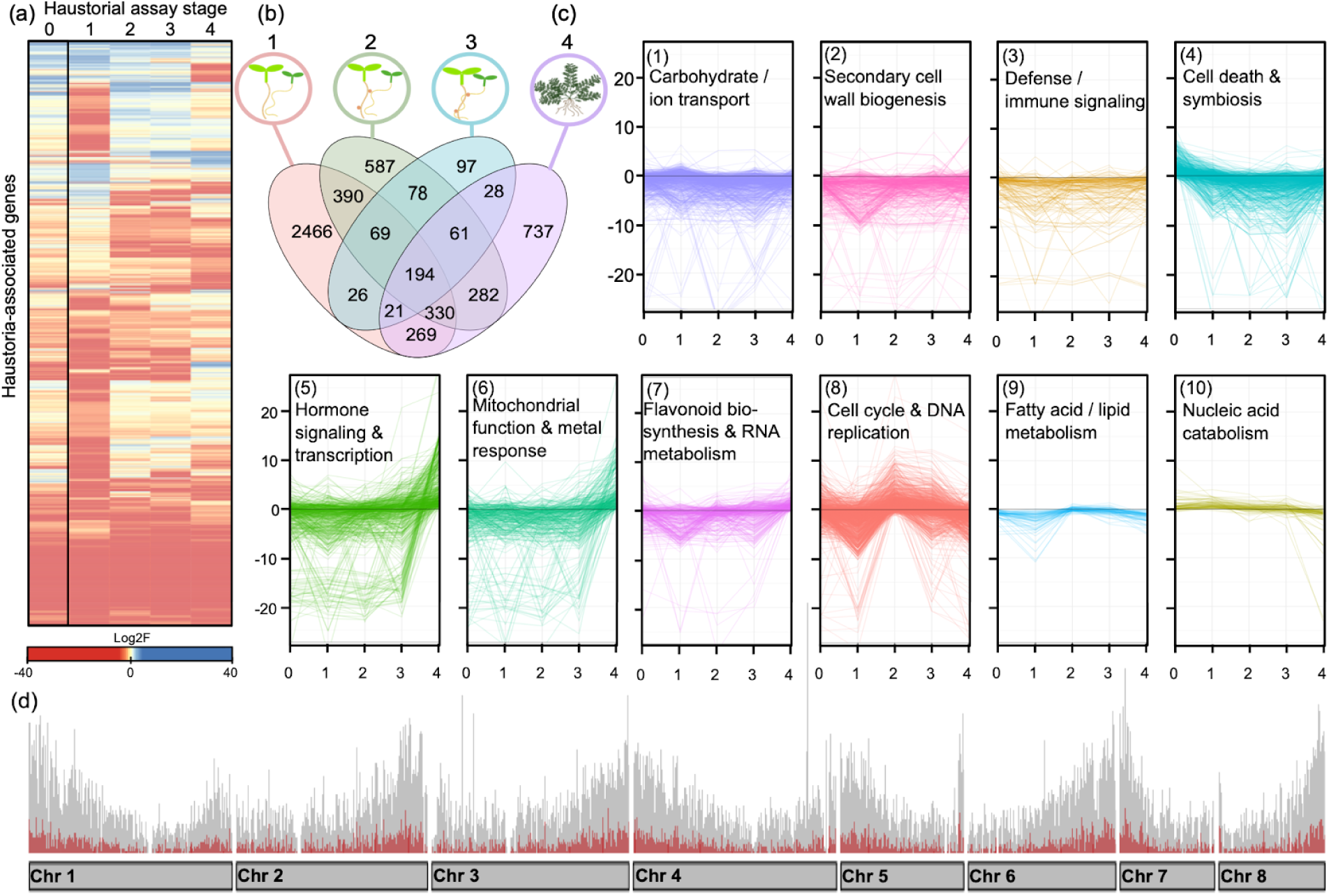
Haustoria-associated genes and their gene expression in *Pedicularis groenlandica*. We identified 5,635 haustoria-associated genes, including both up- and down-regulated genes. **a)** Heatmap of Log2FoldChange values for each haustoria-associated gene. Red indicates down-regulation and blue up-regulation. We include “Stage 0”, young roots pre-host exposure, to show changes in gene expression across the entirety of haustorial development and attachment. **b)** Venn diagram showing the overlap in genes involved in haustorial response across the four stages. Stage 1, response 24 hours post-host exposure, had the highest number of stage-specific genes. **c)** Changes in gene expression among the sets of genes within 10 haustoria-associated WGCNA modules, representing co-expressed molecular responses to host presence, haustorial initiation, and haustorial attachment. Lines in each module plot show the change in gene expression of a single gene throughout the stages of haustorial development. Each module is labeled according to the most enriched GO biological process terms for the set of genes. **d)** Bar chart depicting haustoria-associated gene density (red) compared to general gene density across the genome (grey), calculated in 10kb sliding windows. The grey bars along the horizontal axis represent *P. groenlandica*’s eight chromosomes.

In stage 1, twenty-four hours after host exposure, 3,734 genes were differentially expressed, and showed enrichment for biological process GO terms relating to mitotic cell division, DNA replication, and cell wall biogenesis. In stage 2, when haustoria began to swell, 2,054 differentially expressed genes showed functions enriched in pigment biosynthesis, tetrapyrrole metabolism, ion transport, and unidimensional cell growth. In stage 3, during host invasion, 679 genes were differentially expressed, which were enriched for pollen tube growth, pollination, cellular tip growth, and unidimensional cell growth. In mature haustoria (stage 4), 2,047 genes were differentially expressed, showing enrichment for light response and photosynthesis.

Of the 32 co-expression network modules identified by WGCNA, we identified ten that are significantly correlated (P < 0.05) with haustoria, and assigned functional descriptions to each based on the most enriched GO terms among genes in the module (**Fig. 3c**). These one of the haustoria-associated modules was enriched for pollen tube-associated functions (biological process GO terms like, pollination, unidimensional cell growth, pollen tube development, and pollen tube growth, among others; **Fig. S2**). Conservatively, we focus on the modules identified as significantly associated with haustoria compared to all other tissue-types included in the study (**Fig 3c**).

### Haustoria-associated gene and regulatory motif overlap with non-parasitic tissues

Our set of haustoria-associated genes has the highest overlap with genes associated with pollen tubes; more than any other tissue-type, including roots (**Fig. 4a**). Here, gene overlap means that the same loci were identified as differentially expressed in both haustoria and a particular tissue. Similarly, regulatory motifs enriched in the flanking regions of haustoria-associated genes within individual WGCNA co-expression modules correlated with haustorial tissues showed the highest level of overlap with pollen tubes (**Fig. 4b)**, although the degree of overlap was lower when measured by motif identity (**Fig 4a**). Despite sharing a substantial set of genes, haustoria and pollen tubes often exhibit opposite differential expression patterns: genes up-regulated in pollen tubes are frequently down-regulated in haustoria (**Fig. 4b**). This resulted in the expression profiles of haustoria and pollen tubes being the most distinct from one another among all tissues considered (**Fig. S3**). Finally, across all co-expression modules identified as haustoria-associated during haustoria/pollen tube overlap analyses, haustoria- and pollen tube-associated genes were grouped together more often than we would expect due to chance alone (**Fig 4c**, FDR-adjusted *P*<0.05).

**Fig. 4.**
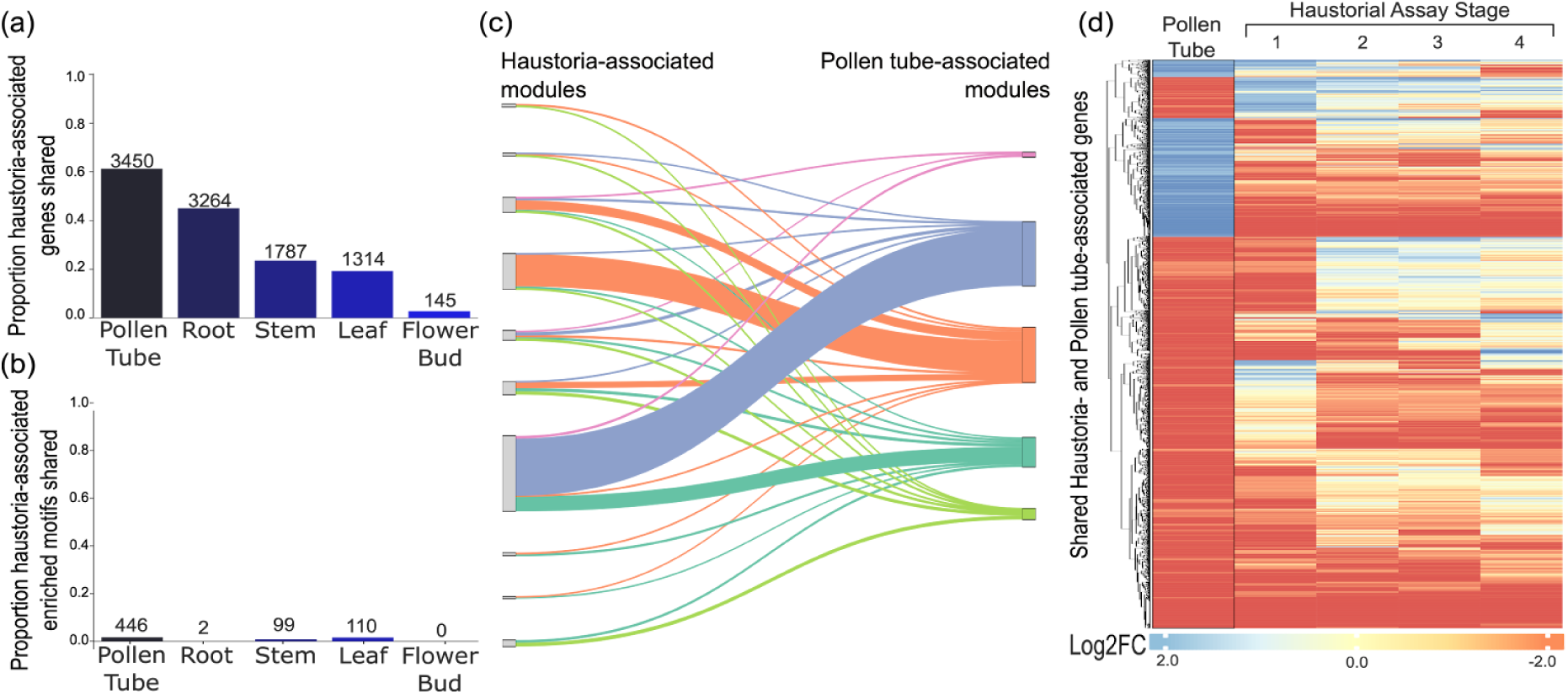
Overlap of haustoria-associated genes and regulatory motifs with non-parasitic tissues. Bar charts showing **a)** the proportion of haustoria-associated genes and **b)** enriched regulatory motifs that overlap with non-parasitic tissues. We additionally report the count of overlapping genes and motifs above each bar. We show that haustoria-associated genes and motifs overlap most often with pollen tube’s. **c)** Relative grouping of genes into co-expression modules in haustoria and pollen tubes. Haustoria- and pollen-associated genes were grouped together in co-expression modules more often than expected due to chance alone (*P*<0.05 in each haustoria-associated module). **d)** Heatmap of gene expression (Log2FoldChange) of overlapping haustoria- and pollen tube-associated genes. Blue values are up-regulated genes, and red are down-regulated genes. Many shared genes show the opposite direction of expression in pollen tubes versus haustoria.

### Divergence between haustoria-associated genes and their paralogs

The 5,635 haustoria-associated genes identified in *P. groenlandica* were grouped into 3,761 orthogroups, including alternative splice isoforms predicted by BRAKER3. After collapsing isoforms (which grouped into the same orthogroup 99.99% of the time), this dataset contains 4,718 *P. groenlandica* genes, corresponding to a mean +/- stdev of 1.25 +/- 0.73 genes per orthogroup.

We identified 2,054 haustoria-associated genes (44%) that have at least one non-haustoria-associated paralog. Of these, 1211 (59%) have a paralog that arose from a duplication after the divergence between Orobanchaceae and *Mimulus*, while 1121 (55%) have a paralog that arose ancestral to this split. Only 278 (14%) of haustoria-associated genes have both types of paralogs in the same gene tree.

We examined the divergence and overlap in tissue-specific expression between paralogs in each category, which, for convenience, we hereafter refer to as paralogs that arose from “recent” versus “ancient” duplications. For genes that were the product of recent duplications, paralogs of haustoria-associated genes were more frequently differentially expressed in pollen tubes (40%) than any other tissue type (3-13%), and the remaining half (52%) were not differentially expressed in any non-parasitic tissue considered (**Fig. 5a**). For ancient paralogs, an even greater proportion are associated with pollen tubes (57%) and very few in roots (19%; **Fig. 5b**). In addition, paralogs expressed in pollen tubes are often exclusively found in this tissue, while root associated paralogs tend to be expressed in multiple tissue-types (**Fig. 5a-b**).

**Fig 5.**
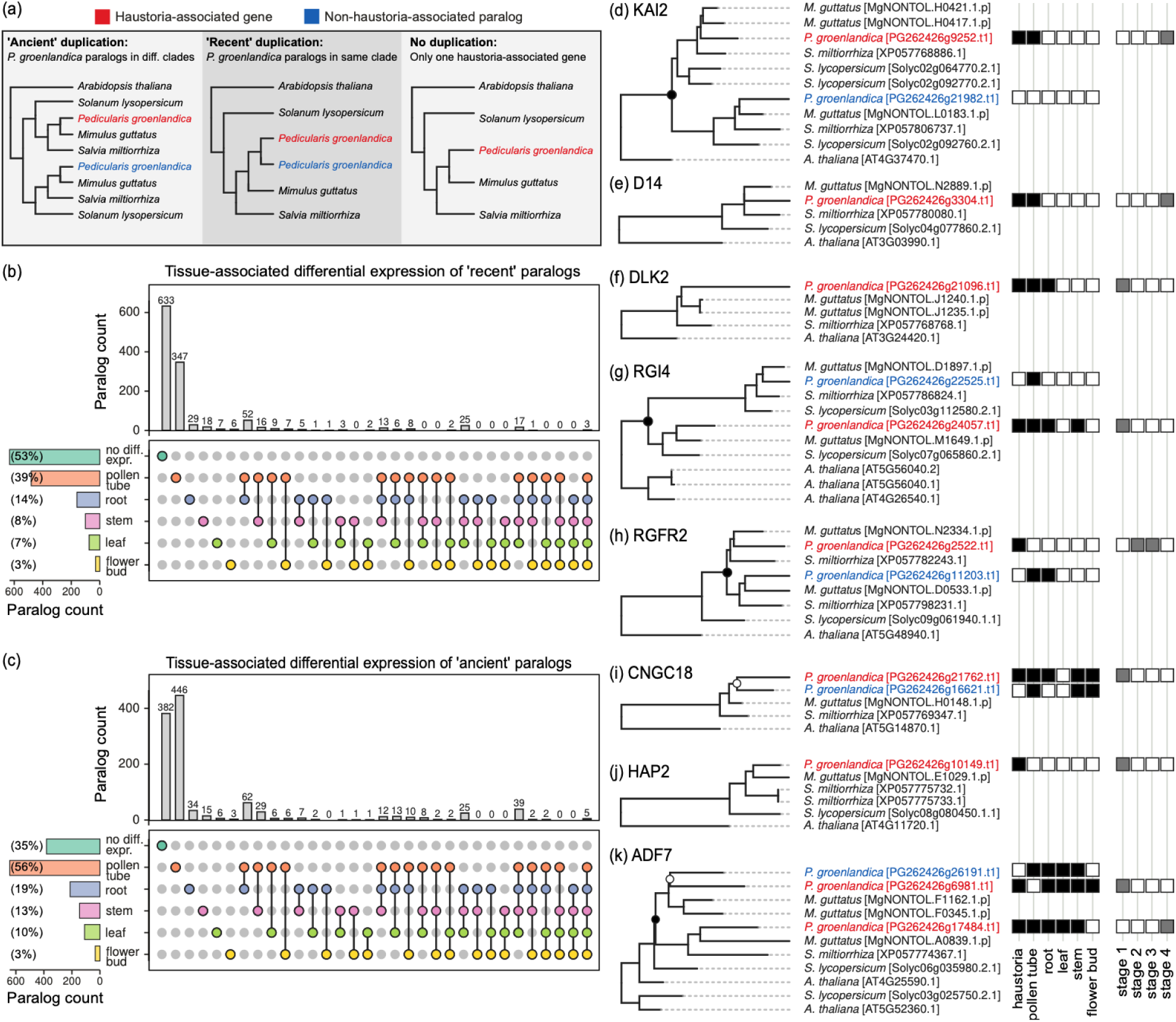
Differential expression of Pedicularis groenlandica haustoria-associated genes and their paralogs across multiple plant tissues. (a) The closest non-haustoria-associated paralog separated by either a “recent” or “ancient” gene duplication was identified in each orthogroup gene tree, if present. (b-c) The expression profiles of non-haustoria-associated paralogs are summarized in upset plots for “recent” (b) and “ancient” (c) paralogs, respectively. Most non-haustoria-associated paralogs of haustoria-associated genes are either not differentially expressed in any tissues, or are differentially expressed in pollen tubes. (d-k) Example orthogroup gene trees showing expression profiles for haustoria-associated genes and their paralogs, if present. These orthogroups contain homologs associated with haustorial development or pollen tube guidance. Open circles indicate “recent” gene duplications, and closed circles “ancient” gene duplications. For haustoria-associated genes we show the haustorial assay stage in which they were differentially expressed.and whether this can be predicted by the presence of recent and/or ancient gene duplications. Using a GEE logistic model, we found that the presence of a recent non-haustorial paralog

### Selected gene trees

We inspected many gene trees and selected several that demonstrate interesting patterns among genes that contain orthologs with demonstrated function in haustorial development and attachment in other parasitic lineages. We detected haustoria-associated genes in both KAI2 and D14, which are receptors of strigolactones in Orobanchaceae used to detect host plants (Mutuku et al. 2020). A haustoria-associated gene was also found within the homolog DLK2, which has not been previously implicated in parasitism. In all three groups the haustoria-associated genes were also differentially expressed in pollen tubes, as well as roots in the case of DLK2. Only KAI2 contains a paralog, which was not differentially expressed in haustoria or any other tissues examined.

Two homologs of ROOT GROWTH FACTOR that are associated with haustorial morphogenesis contain both haustoria-associated genes and non-haustoria-associated paralogs in *P. groenlandica* (**Fig. 5g-h**). In RGI4 an ancient duplication separates a haustoria-associated gene that is differentially expressed in several tissues, including during stage 1 of the haustorial assay, from a non-haustoria-associated paralog that is uniquely expressed in pollen tubes. RGFR2 similarly contains an ancient duplication separating a haustoria-specific gene that was differentially expressed during stage 2 of the haustorial assay, from a paralog that is differentially expressed in pollen tubes and roots.

Three orthogroups were selected that contain haustoria-associated genes among homologs of *Arabidopsis* genes with known function in pollen-tube growth and guidance (CNGC18, HAP2, and ADF7; **Fig. 5i-k**). In CNGC18 and ADF7, we observe a haustoria-associated gene separated by a “recent” duplication from a non-haustoria-associated paralog that is differentially expressed in pollen-tubes and other tissues. In HAP2 the *P. groenlandica* ortholog is haustoria-specific. A haustoria-associated gene in all three orthogroups was expressed during stage 1 of the haustorial assay. Overall, this set of tree visualizations (**Fig. 5d-k**) demonstrates the diversity of patterns of gene duplication and expression divergence across gene families containing haustoria-associated genes in *P. groenlandica*.

### Modeling the effect of gene duplications

To quantify the relative importance of gene duplications for the evolution of haustoria-specific expression, we compared the number of genes that are haustoria-specific to those that are differentially expressed in multiple tissues, increases the odds that a haustoria-associated gene is haustoria-specific (OR=1.59, 95% CI 1.27–2.00, p<0.001). In contrast, ancient non-haustoria-associated paralogs showed no significant association (OR=0.86, 95% CI 0.70–1.05, p=0.13). A direct Wald test confirmed the difference between recent and ancient effects (χ²=23.19, p=1.5×10^-6^). The average marginal effects from the GEE indicate that recent non-haustoria-associated paralogs increase the probability of haustoria-specific expression by 7.6 percent (SE=1.9, p<0.001), whereas ancient paralogs are associated with a non-significant −2.5 percent change (SE=1.7, p=0.13).

### Diverging haustorial gene repertoires across parasitic plant lineages

Among the haustorial transcriptome datasets from five parasitic species (4 Orobanchaceae, 1 *Cuscuta*), we identified a highly variable number of haustorial-associated genes. It was lowest among the two oldest datasets (*S. hermonthica*=120, *O. aegyptiaca*=472). By contrast, a much greater number of haustoria-associated genes was detected in the three recent datasets (*C. campestris*=10,396, *P. japonicum*=1,058, and *P. groenlandica=*5,635). We restricted our main comparative analysis to these three datasets (**Fig. 6a)**. Genes in shared orthogroups were expressed in each of the four stages of haustorial development (**Fig. 6c**). Orthology analysis identified haustoria-associated gene families shared between Orobanchaceae and *Cuscuta*, and an even greater number that are lineage-specific (**Fig. 6b**). Within the five species dataset, we identified only two orthogroups shared among all Orobanchaceae, compared to a majority of species-specific orthogroups (**Fig. S3**). The highest orthogroup overlap was among the three modern datasets, with 660 shared between *P. groenlandica* and *C. campestris*, 258 shared between *P. japonicum* and *P. groenlandica*, and 90 shared between *C. campestris* and *P. japonicum*. Among all five species, the highest number of overlapping orthogroups occurred in those that contained genes which were differentially expressed during haustorial swelling (stage 2).

**Fig. 6.**
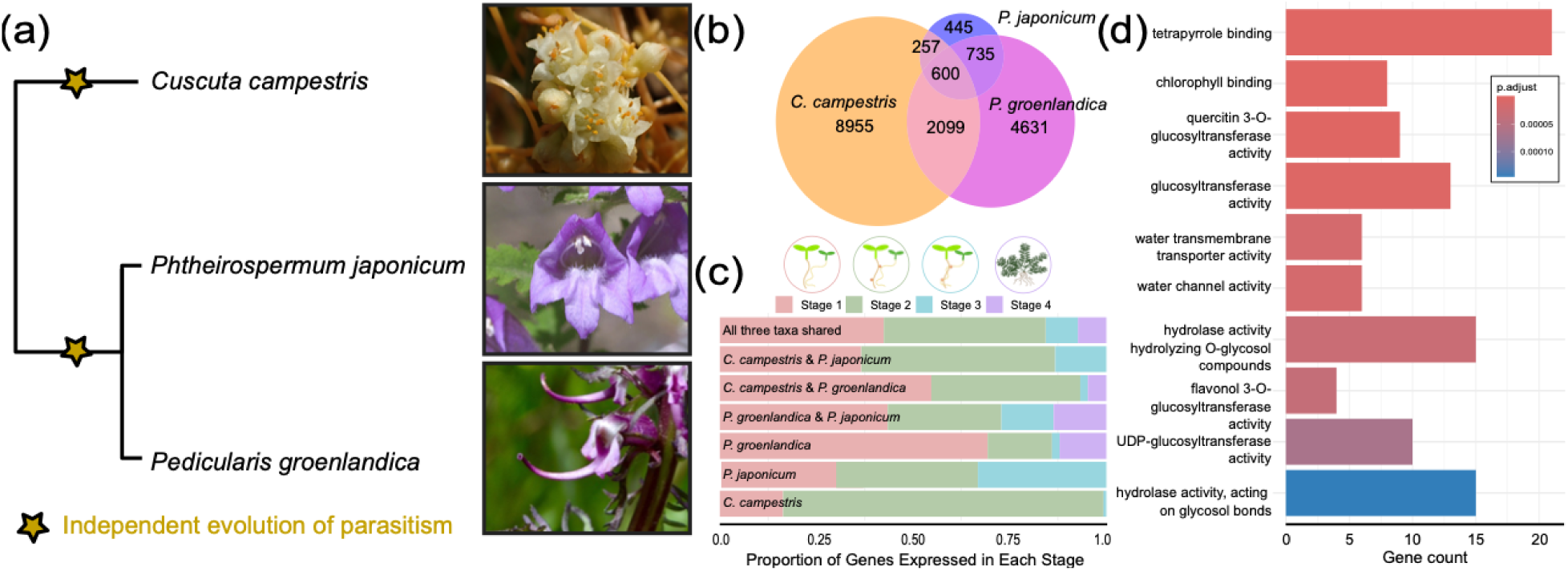
Orthology of haustorial-associated genes in *Pedicularis groenlandica*, *Phtheirospermum japonicum*, and *Cuscuta campestris*. **a)** Cladogram showing the relationships between the three species, with two independent transitions to parasitism. Photographs of *C. campestris* and *P. japonicum* taken from iNaturalist with permission. (https://www.inaturalist.org/photos/62167474; https://www.inaturalist.org/observations/245056018) **b)** Venn diagram of the overlap in orthogroups among the three species. The number in each section is the sum of genes in all three species that make-up orthogroups in each category. **c)** Bar chart depicts the breakdown of genes in each of the Venn diagram categories by haustorial assay stage (1-4) in which they were differentially expressed in each species. *C. campestris* has a high amount of species-specific stage 2 genes. **d)** Molecular function GO enrichment analysis of the 600 genes in the orthogroups that overlap among all three species. Color of bars shows the adjusted p-value associated with enrichment, with red being more enriched and blue less enriched.

We evaluated the number of genes that made up the orthogroups in the three modern datasets. There were 600 haustoria-associated genes in the orthogroups that were shared by all three species: *P. groenlandica*, *P. japonicum*, and *C. campestris* (**Fig. 6b**). These genes included those involved in tetrapyrrole and chlorophyll binding, aquaporins, glucosyltransferases, hydrolases, and flavonoid-modifying enzymes (**Fig. 6d**). The lineage-specific orthogroups contained genes with distinct but complementary functions. In Orobanchaceae (overlap between *P. groenlandica* and *P. japonicum*), genes were enriched for functions related to cell wall modification, tropism, and photosynthesis/oxidative stress. In *C. campestris*, a holoparasite, genes were enriched for functions like external encapsulating structure organization, cell wall organization and modification, pectin catabolic process, and pattern specification.

## Discussion

### Pedicularis groenlandica genome

Our *P. groenlandica* genome is a high-quality, chromosome-scale addition to the growing body of molecular data in the parasitic plant family Orobanchaceae. Including *P. groenlandica*, there are now eleven publicly-available chromosome-scale genomes in the family (Ma *et al*., 2021; Xu *et al*., 2022; Twyford *et al*., 2024; Kim *et al*., 2024; Chen *et al*., 2024; Bürger *et al*., 2025), and three in *Pedicularis* alone (Xu *et al*., 2022; Fang *et al*. 2024). HiC data supported scaffolding the genome into eight chromosomes (**Fig. 2d**), which was consistent with our expectations based on available chromosome count data in *P. groenlandica* (Löve, 1982) and across *Pedicularis* (IPCN chromosome reports), including two chromosome-scale genomes (Xu et al., 2022; Fang et al., 2024). We report relatively constant gene density and GC content across the eight chromosomes and high transposable element (TE) density (**Fig. 2d**).

The *P. groenlandica* genome is highly repetitive (76.5% repeated), of which 44.5% were annotated as transposable elements (TE’s). Differential accumulation of TE’s drives changes in genome size in plants, and in parasitic plants in particular (Neumann *et al*., 2021). Parasitic plants may disproportionately accumulate TE’s relative to non-parasitic plants because genes “hitchhiking” on TE’s can be horizontally transferred into parasitic plant genomes, driving genetic innovation, even in the face of increased rates of gene loss following the transition from autotrophy to heterotrophy (Cai, 2023). *P. groenlandica* has similarly high repeat content to other parasitic Orobanchaceae (*Phelipanche aegypitiaca*=84.4%, *Orobanche cumana*=79.8%, *Orobanche coreulescens*=86.3%), and higher repeat content than non-parasitic Orobanchaceae (*Lindenbergia luchunensis*=30.7%, *Rehmannia glutinosa*=67%; Ma *et al*. 2021; Kim *et al*., 2024). The relationship of genome repeat content to parasitism warrants further exploration, particularly in a comparative context.

Xu et al. (2022) conducted a comparative genomic study of Orobanchaceae genomes, which included the *P. cranolopha* genome in some analyses. They observed limited synteny between *P. cranolopha* and two other Orobanchaceae genomes (*Lindenbergia luchunensis* and *Orobanche cumana var. cumana*). Consistent with Xu et al.’s (2022) findings, *Pedicularis groenlandica*’s eight chromosomes show clear synteny with *P. cranolopha*, with some small inversions and translocations, but a high amount of rearrangement with respect to *O. cumana* (**Fig. 2b**). From this, we conclude that *Pedicularis* genomes have undergone a major re-organization since splitting from the common Orobanchaceae ancestor.

### Haustorial gene function and regulation in *P. groenlandica*

Haustoria-associated genes make-up 20.65% of all protein-coding genes in *P. groenlandica*, indicating haustorial development and attachment leverages a wide range of molecular processes and elicits a genome-wide response. Functions of genes across the haustorial developmental stages (**Fig. 4b**) were consistent with the functional categories we expected to be related to parasitism based on previous haustorial RNAseq studies. For example, during Stage 1, we observed the differential expression of genes involved in cell division and cell wall expansion, which likely is related to the initiation of haustorial growth (Ishida *et al*., 2016; Cui *et al*., 2020). Stages 3 and 4 had similar expression patterns to each other, both showing enrichment for cellular tip growth-related genes. Cui et al. (2016) found that cellular tip growth and cell elongation are major mechanisms by which haustoria and functional haustorial hairs form in *Phtheirospermum japonicum* (Orobanchaceae), a facultative hemi-parasite that is closely related to *Pedicularis* within Orobanchaceae.

Similarly, the functional associations of haustoria-associated gene co-expression networks included biological processes with known relevance to haustorial growth and invasion (**Fig. 3c**). For example, flavonoid biosynthesis. Flavonoids are known to be related to haustoria formation in *Triphysaria versicolor* (Albrecht *et al*., 1999) and in the genus *Cuscuta* (Lozanova *et al*., 2025). We additionally identified a module of genes related to defense and immune response (**Fig. 3c**), which we expect to include genes interacting with host defense pathways during parasitic invasion. Hormone signaling and ion transport may also relate to host-parasite cross-talk, and we saw enrichment of genes involved in auxin, ethylene, salicylic acid, and phenylpropanoids, each of which have been linked to host-parasite signaling in other parasitic plants (Ishida *et al*., 2016; Cui *et al*., 2020; Wakatake *et al*., 2020; Mutuku *et al*., 2021). Our results thus tell a consistent functional story about haustorial development compared to other parasitic lineages in which haustoria-associated genes have been identified, and we are confident that we have successfully captured the haustorial developmental process in *P. groenlandica*.

### Pollen tubes and the evolution of haustoria-associated genes

Both Yang et al. (2015) and Yoshida et al. (2019) found that homologs of haustoria-associated genes in *Tryphisaria versicolor*, *Phelipanche aegypotiaca*, *Striga hermonthica*, and *S. asiatica* were enriched in pollen-related functions in model species like *Arabidopsis thaliana* and *Zea mays*. We hypothesized that this pattern results from the overlap in function of haustoria and the process of pollen tube growth which, like haustorial invasion, requires signal detection, growth through foreign tissue, and directed unidimensional cell growth. Sixty-one percent (3450/5635) of haustoria-associated genes were differentially expressed in both haustoria and pollen tubes, which represented a significant overlap (Fischer’s Exact Test P<10^-70^). This overlap was greater than any other non-parasitic tissue considered, including roots (**Fig. 4a**). The extent of the haustoria-pollen tube relationship was surprising, as haustoria-associated genes have been previously hypothesized to be co-opted from predominantly root-associated genetic pathways.

Pollen tube growth is known to require ethylene and auxin signaling (Guan *et al*., 2013; Chen *et al*., 2014; Weijers & Wagner, 2016; Cui *et al*., 2020; Mou *et al*., 2020) as well as genes in the expansin superfamily (Cosgrove, 1997; Sampedro & Cosgrove, 2005), all of which are also correlated with haustoria development (Yoshida *et al*., 2016, 2019). In addition to high overlap in differential expressed genes among haustoria and pollen tubes, the two tissues shared the highest number of enriched regulatory motifs. The most common transcription factor family that shared enriched motifs interact with was ERF/DREB (**Fig. S4b**), which regulates ethylene signaling. Yoshida *et al*. (2016) argue that the co-option of the lateral root development program led to the evolution of haustoria across the twelve parasitic plant lineages, focusing in particular on pathways involved in auxin-biosynthesis and regulation. Across 5,635 haustoria-associated genes in *P. groenlandica*, 155 were involved in known auxin pathways, and of those, 62 auxin genes were additionally associated with pollen tubes, 18 were associated with roots, and 33 were associated with all three tissues. Despite known functioning of auxin pathways in roots, our results suggest that, in *P. groenlandica*, auxin functionality in haustoria has predominantly been co-opted from pollen tube-associated auxin pathways.

One explanation for the high proportion of haustoria-associated genes that were differentially expressed in pollen tubes is that genes function pleiotropically in the two tissues. We expect pleiotropy to give rise to divergence in *cis*-regulatory elements that mediate gene expression, which is supported by our observation that a small proportion of enriched regulatory motifs are shared between haustoria and non-parasitic tissues compared to protein-coding loci (**Fig. 4b**). Further, genes being similarly grouped into coexpression modules (**Fig. 4c**) and disparity in the direction of expression (**Fig. 4d**) among shared haustoria- and pollen-tube associated genes is consistent with pleiotropy, and suggests that this pleiotropy may be antagonistic in haustoria and pollen tubes (*i.e.* mutations in a pleiotropically functioning gene may be beneficial in one tissue, but deleterious in another; Hoekstra & Coyne, 2007). Together, these results suggest that divergence in regulation (*i.e.* gene expression) mediates the variability in the functional processes of haustoria and pollen tubes.

A majority of the 5,635 haustoria-associated genes in *P. groenlandica* showed evidence of association with pollen tubes, either because they were differentially expressed in pollen tubes, nested within pollen tube-associated gene families, or both. Among haustoria-associated genes that did not group into an orthogroup with a *P. groenlandica* paralog, 66% (1763/2664) were themselves differentially expressed in pollen tubes; among those that did contain a paralog in their orthogroup, 52% (1061/2054) were members of pollen tube-associated gene families (*i.e.* at least one paralog was differentially expressed in pollen tubes). These patterns suggest that haustoria have disproportionately co-opted functions from pollen tubes, and that pollen tube-associated genes and gene families have been integral to the evolution of haustoria-associated genes in *Pedicularis*. Although shared function cannot be inferred from expression alone, our findings provide strong support for a close relationship between these two tissue types that future studies can address experimentally to test pleiotropy more directly.

### Alternative explanations for the co-option of haustoria-associated genes

Yang et al. (2015) argue that haustoria-associated genes evolve through regulatory neofunctionalization following ancient gene duplications, which they correlate with ancestral WGD’s. If haustoria-associated genes were derived from WGD’s, we expected to identify an ancient duplication in the evolutionary history of the majority of haustoria-associated genes in *P. groenlandica* and likely haustoria-specific gene expression. In *P. groenlandica*, only 19.2% of haustoria-associated genes showed evidence of diverging from non-haustoria-associated paralogs following an ancient duplication. Furthermore, from examination of gene tree topologies alone, the methodology employed by both Yang et al. (2015) and this study, whether ancient duplications were derived from a WGD or another type of gene duplication (*i.e.* a tandem duplication) that occurred in a shared common ancestor cannot be distinguished. We show that the evolution of haustoria-specific expression is significantly more likely following a recent duplication, and is not affected by the presence of ancient duplications. This is consistent with the idea that recent, rather than ancient, duplications have facilitated neofunctionalization of haustoria-specific genes. The predictive effect, however, is quite small, as the presence of a recent paralog only makes it 7.6% more likely that a gene will exhibit haustoria-specific gene expression.

Both regulatory neofunctionalization and pleiotropy are contexts in which new phenotypes can evolve through variation in gene expression. Within our gene tree dataset, we observed topologies and differential expression patterns in haustoria-associated and non-haustoria-associated paralogs consistent with three, non-mutually exclusive, evolutionary explanations for the advent of haustorial function: (1) pleiotropy (2) neofunctionalization following a recent duplication, and (3) neofunctionalization following an ancient duplication. We hypothesize that co-option of non-parasitic functions for haustorial development may initially occur via pleiotropy, and subsequent recent (potentially lineage-specific) gene duplications have given rise to paralogs primarily through neofunctionalization, perhaps to alleviate the burden of pleiotropy. Fishman *et al*. (2025) provide evidence of neofunctionalization following a recent, segmental tandem duplication in RGF orthologs in *Phtheirospermum japonicum*, consistent with our hypothesis for haustoria-associated gene evolution. Further studies that validate haustoria-associated gene functions in non-parasitic tissues and compare patterns of selection acting on genes uniquely versus non-uniquely expressed in haustoria will be needed to comprehensively investigate this hypothesis.

### Divergent pathways towards the evolution of haustoria

We expected to identify a core set of genes across Orobanchaceae that contribute to haustorial function in the family, derived in Orobanchaceae’s most recent common ancestor. Instead, although there is some overlap among haustoria-associated genes across the parasitic plant lineages considered, a high proportion of haustoria-associated genes are species-specific (**Fig. 6**). Yang et al. (2015) similarly found that most haustoria-associated genes were lineage-specific when comparing *Triphysaria versicolor, Striga hermonthica* and *Phelipanche aegyptiaca*. Unlike our analyses, the species they examined span the full spectrum of parasitism, including a facultative hemiparasite, obligate hemiparasite, and obligate holoparasite. Their results suggest that hemiparasitic vs. holoparasitic haustorial function necessitates the recruitment of diverse sets of genes, with relatively small overlap.

It was therefore surprising that even between the two closely-related facultative hemiparasites *P. groenlandica* and *P. japonicum*, a high proportion of haustoria-associated genes were species-specific. This result is particularly interesting in light of our findings that 1) *Pedicularis* have highly rearranged genomes with respect to other Orobanchaceae and 2) recent duplications predict whether a haustoria-associated paralog is differentially expressed only in haustoria. Together, these results suggest that since the evolution of haustoria in the common ancestor of Orobanchaceae, a substantial amount of variation has evolved among descendant lineages in the recruitment of genes into haustoria-associated functions. Many aspects of the biology of these organisms have likely impacted this process. For example, *Pedicularis* has experienced an adaptive radiation since the Miocene, where it now comprises >600 species distributed mostly throughout montane and arctic ecosystems (Eaton & Ree, 2013). Whether *Pedicularis* has evolved variation in haustoria-associated regulatory neofunctionalization faster than other parasitic lineages will require further comparative analyses, examining the distribution of haustoria-associated genes with respect to synteny. The closely related genus *Castilleja* offers an opportunity for comparison with a similarly large and diverse clade (Li *et al*., 2019).

Because transcriptomes of *Cuscuta campestris* haustoria were sequenced more recently (Bawin *et al*., 2022, 2024), neither Westwood et al. (2012) nor Yang et al. (2015) included a convergently evolved haustoria from outside Orobanchaceae in their comparative transcriptomic analyses. Using Bawin et al.’s (2024) haustoria RNAseq data, we were able to compare haustoria-associated genes in Orobanchaceae and *Cuscuta*. There were a number of haustorial genes that occurred in common orthogroups among *P. groenlandica*, *P. japonicum*, and *C. campestris*. The shared orthogroups among all three lineages included genes with functions related largely to water transfer. Both *P. groenlandica* and *P. japonicum* are hemiparasitic, meaning their haustoria invade only the xylem of a host plant in order to siphon water (Yoshida *et al*., 2016). In contrast, *C. campestris*, as a holoparasite, has haustoria that penetrate both the xylem and phloem of a host (Yoshida *et al*., 2016). The overlap in genes that function in tetrapyrrole binding, glucosyltransferase, and water transmembrane/channel molecular pathways (**Fig. 6D**), indicates potential commonality in the way that hemi- and holoparasites invade xylem and transfer water from their hosts. This is consistent with the hypothesis that holoparasitism evolves subsequent to the evolution of hemiparasitism, and that the first haustoria in parasitic lineages likely resembled the lateral haustoria of hemiparasites (Westwood et al. 2010).

## Conclusions

This study presents strong evidence for the importance of pollen tube-associated genetic pathways in the evolution of haustoria in *P. groenlandica*. We additionally highlight the need to evaluate diverse mechanistic explanations for the co-option of genes for haustorial function in parasitic plants, and, more broadly, the ways in which changes in gene expression can contribute to the evolution of novel phenotypes. Finally, although the data available from the Parasitic Plant Genome Project represented a huge advance in parasitic plant genetic research, these data no longer meet the expectations for modern RNA sequencing experiments, including the expected level of replication (three biological replicates per stage), that is available in the *P. groenlandica*, *P. japonicum*, and *C. campestris* datasets. We underscore the need for additional high-quality genome and haustorial RNA datasets in order to conduct more comprehensive comparative studies of haustorial gene evolution within Orobanchaceae and across parasitic plant lineages.

## Acknowledgements

We thank B. Stokes for her help with tissue collection at RMBL, and A. Cisse for assistance with both fieldwork and seed germination for haustorial assays. We additionally thank N. Gershberg for his assistance and advice related to plant care, and J. Reithel and R. Williams for their assistance in fieldwork at RMBL. We thank I. Overcast, J. Leung, Y. Yang, and R. Rampalli, and D. Timerman for their feedback on the manuscript.

This work was supported by the National Science Foundation (NSF DEB-2046813) awarded to D.E and, the RMBL Graduate Student Grant awarded to R.O.C., and the Botanical Society of America Graduate Student Research Award awarded to R.O.C.

## Author contributions

R.O.C and D.E. contributed equally to study design. R.O.C. collected all data and conducted all differential expression analyses. R.O.C. and A.C. did gene co-expression network and motif enrichment analyses. R.O.C. and D.E. conducted the gene duplication-related analyses. R.O.C., D.E., A.C., and L.C. contributed to the writing of the manuscript.

## Data availability

Herbarium vouchers for *P. groenlandica* were deposited at Rocky Mountain Biological Laboratory (RMBL) herbarium (voucher RMBL0007650) and New York Botanical Garden William and Lynda Steere Herbarium. Due to a backlog of vouchers, our specimens had not yet been processed at the time of publication. They will be able to be found using the collector’s and species’ names (R. Cohen) after they have been fully processed by NYBG.

All genomic and transcriptomic data will be available on NCBI, and we will provide the SRAs. Dataframes containing haustorial gene expression and haustoria-associated gene orthology data and the jupyter notebook containing code for gene tree analyses will be available on Dryad.

