## Supplemental Figures for "A reference genome and transcriptome of haustorial development in *Pedicularis groenlandica* reveal diverse trajectories of haustoria-associated gene evolution in parasitic plants"

A1 - PG ☑ Show Statistics

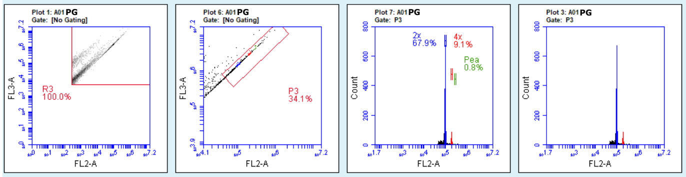


| Plot 1: A01 PG | Count | Events / µL | % of This Plot | % of All | Mean FL2-A | Mean FL3-A | CV FL2-A | CV FL3-A | Median FL2-A | Median FL3-A |
|---|---|---|---|---|---|---|---|---|---|---|
| All | 4,991 | 147 | 100.00% | 100.00% | 40,097.51 | 634,034.50 | 153.28% | 146.95% | | |
| R3 | 4,991 | 147 | 100.00% | 100.00% | 40,097.51 | 634,034.50 | 153.28% | 146.95% | | |

| Plot 6: A01 PG | Count | Events / µL | % of This Plot | % of All | Mean FL2-A | Mean FL3-A | CV FL2-A | CV FL3-A | Median FL2-A | Median FL3-A |
|---|---|---|---|---|---|---|---|---|---|---|
| All | 4,991 | 147 | 100.00% | 100.00% | 40,097.51 | 634,034.50 | 153.28% | 146.95% | | |
| P3 | 1,700 | 50 | 34.06% | 34.06% | 111,158.28 | 1,633,033.97 | 50.87% | 52.48% | | |

| Plot 7: A01 PG Gated on P3 | Count | Events / µL | % of This Plot | % of All | Mean FL2-A | CV FL2-A | Median FL2-A |
|---|---|---|---|---|---|---|---|
| This Plot | 1,700 | 50 | 100.00% | 34.06% | 111,158.28 | 50.87% | |
| 4x (178,336.0 / … | 155 | 5 | 9.12% | 3.11% | 197,683.34 | 4.43% | |
| 2x (90,202.0 / 1… | 1,154 | 34 | 67.88% | 23.12% | 99,056.35 | 4.26% | |
| NullSubset | | | | | | | |
| Pea (260,432.0 /… | 13 | 0 | 0.76% | 0.26% | 284,955.85 | 4.85% | |

| Plot 3: A01 PG Gated on P3 | Count | Events / µL | % of This Plot | % of All | Mean FL2-A | CV FL2-A | Median FL2-A |
|---|---|---|---|---|---|---|---|
| This Plot | 1,700 | 50 | 100.00% | 34.06% | 111,158.28 | 50.87% | |

A2 - PG and pea ☑ Show Statistics

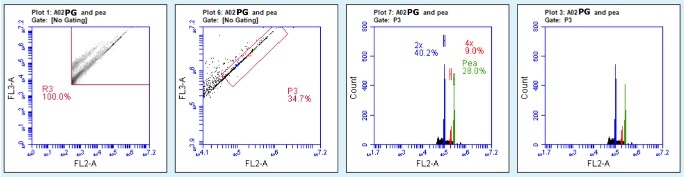


| Plot 1: A02 PG and pea | Count | Events / µL | % of This Plot | % of All | Mean FL2-A | Mean FL3-A | CV FL2-A | CV FL3-A | Median FL2-A | Median FL3-A |
|---|---|---|---|---|---|---|---|---|---|---|
| All | 8,449 | 130 | 100.00% | 100.00% | 64,009.05 | 944,737.73 | 173.87% | 160.24% | | |
| R3 | 8,449 | 130 | 100.00% | 100.00% | 64,009.05 | 944,737.73 | 173.87% | 160.24% | | |

| Plot 6: A02 PG and pea | Count | Events / µL | % of This Plot | % of All | Mean FL2-A | Mean FL3-A | CV FL2-A | CV FL3-A | Median FL2-A | Median FL3-A |
|---|---|---|---|---|---|---|---|---|---|---|
| All | 8,449 | 130 | 100.00% | 100.00% | 64,009.05 | 944,737.73 | 173.87% | 160.24% | | |
| P3 | 2,929 | 45 | 34.67% | 34.67% | 175,984.51 | 2,505,256.53 | 64.25% | 63.10% | | |

| Plot 7: A02 PG and pea Gated on P3 | Count | Events / µL | % of This Plot | % of All | Mean FL2-A | CV FL2-A | Median FL2-A |
|---|---|---|---|---|---|---|---|
| This Plot | 2,929 | 45 | 100.00% | 34.67% | 175,984.51 | 64.25% | |
| 4x (178,336.0 / 223,827.0) | 264 | 4 | 9.01% | 3.12% | 204,928.02 | 5.35% | |
| 2x (90,202.0 / 113,211.0) | 1,178 | 18 | 40.22% | 13.94% | 103,014.22 | 4.71% | |
| Pea (260,432.0 / 326,865.0) | 821 | 13 | 28.03% | 9.72% | 293,895.96 | 4.80% | |

| Plot 3: A02 PG and pea Gated on P3 | Count | Events / µL | % of This Plot | % of All | Mean FL2-A | CV FL2-A | Median FL2-A |
|---|---|---|---|---|---|---|---|
| This Plot | 2,929 | 45 | 100.00% | 34.67% | 175,984.51 | 64.25% | |

A3 - pea ☑ Show Statistics

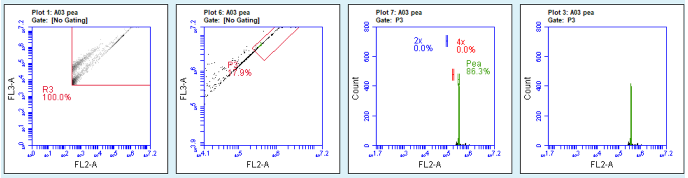


| Plot 1: A03 pea | Count | Events / µL | % of This Plot | % of All | Mean FL2-A | Mean FL3-A | CV FL2-A | CV FL3-A | Median FL2-A | Median FL3-A |
|---|---|---|---|---|---|---|---|---|---|---|
| All | 5,667 | 77 | 100.00% | 100.00% | 86,471.28 | 1,229,885.06 | 198.58% | 184.18% | | |
| R3 | 5,667 | 77 | 100.00% | 100.00% | 86,471.28 | 1,229,885.06 | 198.58% | 184.18% | | |

| Plot 6: A03 pea | Count | Events / µL | % of This Plot | % of All | Mean FL2-A | Mean FL3-A | CV FL2-A | CV FL3-A | Median FL2-A | Median FL3-A |
|---|---|---|---|---|---|---|---|---|---|---|
| All | 5,667 | 77 | 100.00% | 100.00% | 86,471.28 | 1,229,885.06 | 198.58% | 184.18% | | |
| P3 | 1,016 | 14 | 17.93% | 17.93% | 406,826.76 | 5,593,830.13 | 41.34% | 30.97% | | |

| Plot 7: A03 pea Gated on P3 | Count | Events / µL | % of This Plot | % of All | Mean FL2-A | CV FL2-A | Median FL2-A |
|---|---|---|---|---|---|---|---|
| This Plot | 1,016 | 14 | 100.00% | 17.93% | 406,826.76 | 41.34% | |
| 4x (178,336.0 / … | 0 | 0 | 0.00% | 0.00% | 0.00 | 0.00% | |
| 2x (90,202.0 / 1… | 0 | 0 | 0.00% | 0.00% | 0.00 | 0.00% | |
| Pea (326,865.0 … | 877 | 12 | 86.32% | 15.48% | 375,958.22 | 3.72% | |

| Plot 3: A03 pea Gated on P3 | Count | Events / µL | % of This Plot | % of All | Mean FL2-A | CV FL2-A | Median FL2-A |
|---|---|---|---|---|---|---|---|
| This Plot | 1,016 | 14 | 100.00% | 17.93% | 406,826.76 | 41.34% | |

**Fig. S1.** Flow-cytometry report for *Pedicularis groenlandica* (PG) compared to pea.

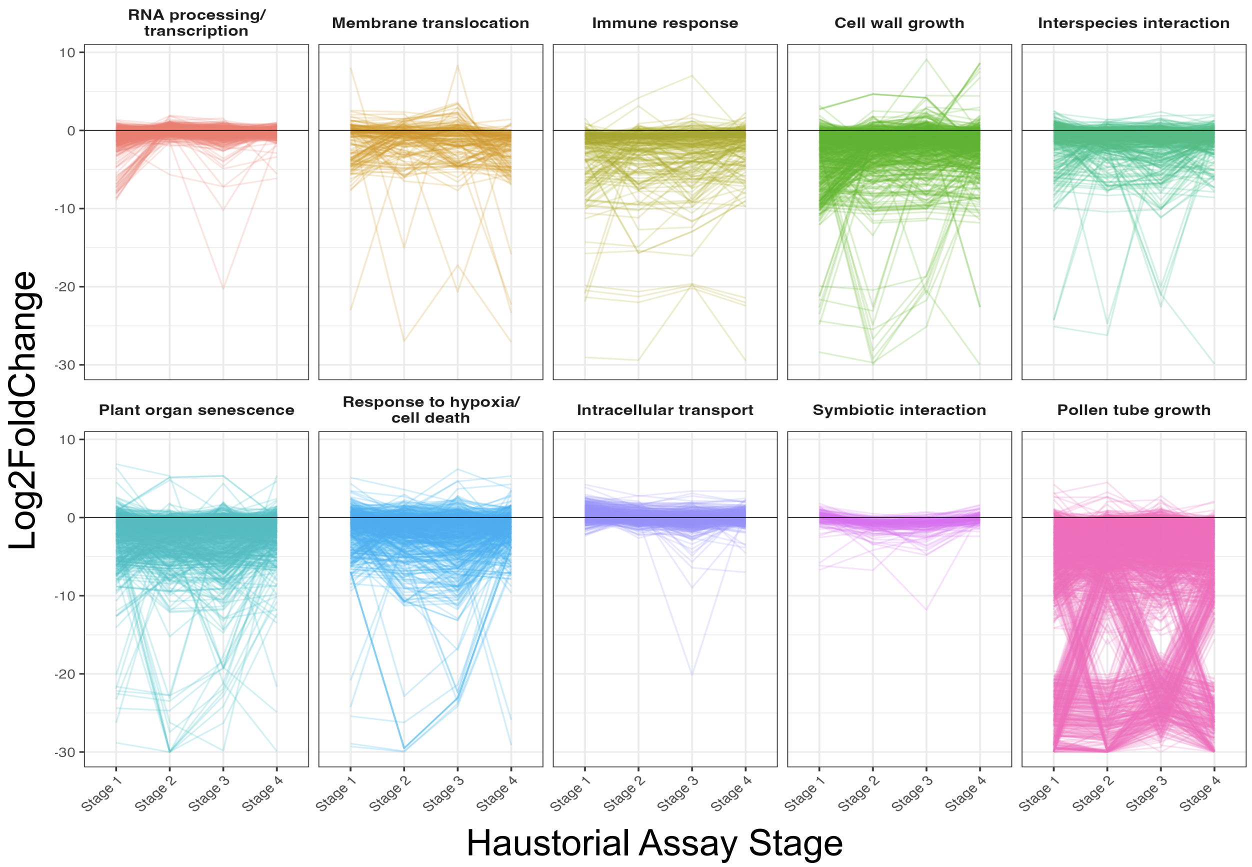


**Fig. S2.** Haustoria-associated WGCNA modules when haustorial tissues are compared to all non-parasitic tissues except pollen tubes. We show change in gene expression (Log2FoldChange) for genes in each module across the four stages of our haustorial growth assay. When pollen tubes are excluded, we find one module with functions specifically related to pollen tube growth.

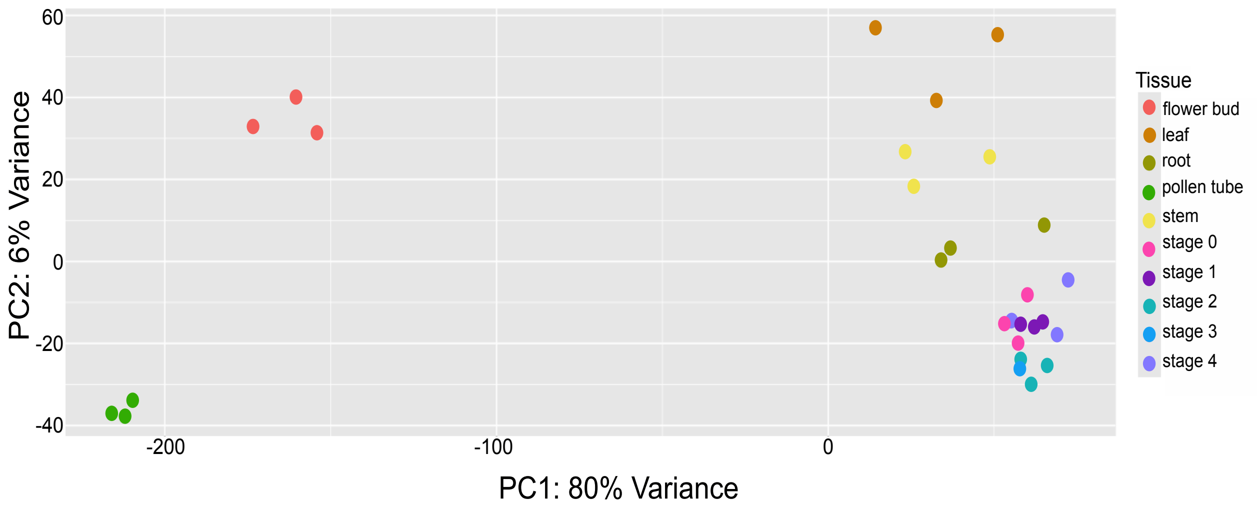


**Fig. S3.** Principal component analysis (PCA) based on differential expression of each tissue, calculated in DESeq2. We show that pollen tubes (bottom left) have distinct differential expression profiles with respect to other tissues, in terms of up- vs. down-regulation. All other tissues group as expected, with haustoria and roots, as well as leaves and stems grouping together.

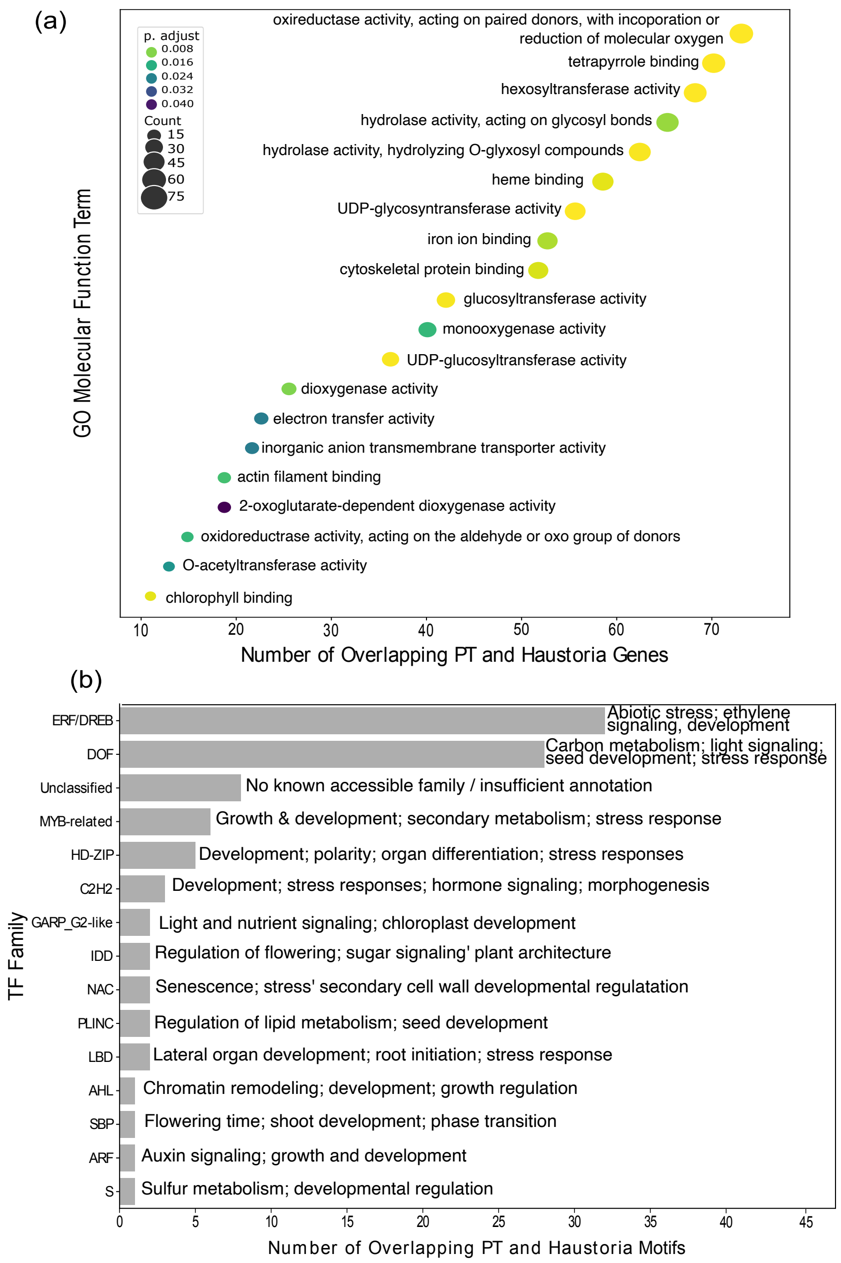


**Fig. S4.** Functional annotations of overlapping haustoria- and pollen-tube associated genes and enriched motifs. **a)** GO term molecular function enrichment analysis of the 3,450 overlapping haustoria-associated and pollen tube-associated genes (PT). Color of the circles shows the adjusted p-value (lighter color/lower value are more significantly enriched). Dot area indicates the number of genes in that GO molecular function category. **b)** Bar chart showing the transcription factor (TF) families overlapping haustoria- and pollen-tube associated regulatory motifs correspond to.

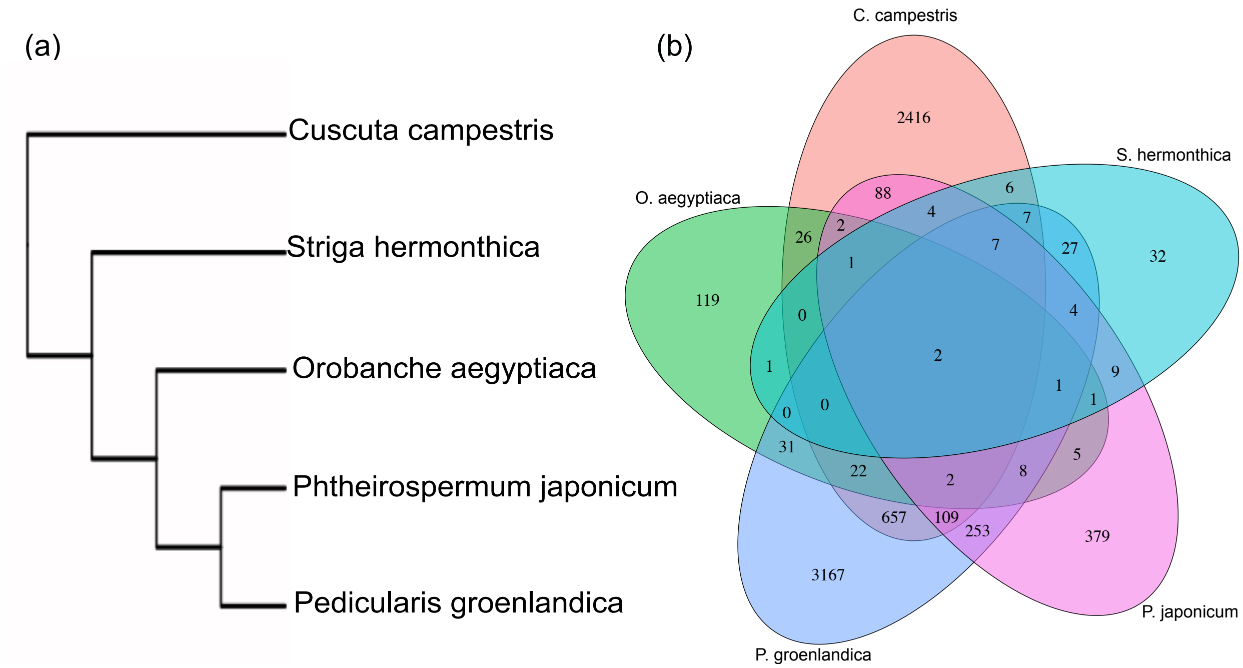


**Fig. S5**. Orthology analysis of haustoria-associated genes across five parasitic plant lineages. **a)** Cladogram of the five species included. Four (*Striga hermonthica*, *Orobanche aegyptiaca*, *Ptheirospermum japonicum*, and *Pedicularis groenlandica*) are in Orobanchaceae, whereas *Cuscuta campestris* represents an independent transition to parasitism. **b)** Venn diagram showing the overlap in orthogroups that contain haustoria-associated genes among the five species. We identified more genes in the three modern datasets (*P. japonicum*, *P. groenlandica*, and *C. campestris*), and they exhibit much higher orthogroup overlap with each other than *S. hermonthica* and *O. aegyptiaca*.
